# Multi-organ analysis of low-level somatic mosaicism reveals stage- and tissue-specific mutational features in human development

**DOI:** 10.1101/2021.08.23.457440

**Authors:** Hyeonju Son, Ja Hye Kim, Il Bin Kim, Myeong-Heui Kim, Nam Suk Sim, Dong-Seok Kim, Junehawk Lee, Jeong Ho Lee, Sangwoo Kim

**Affiliations:** Department of Biomedical Systems Informatics, Graduate School of Medical Science, Brain Korea 21 Project, Yonsei University College of Medicine, Seoul, Republic of Korea; Graduate School of Medical Science and Engineering, Korea Advanced Institute of Science and Technology (KAIST), Daejeon, Republic of Korea; Department of Psychiatry, Hanyang University Guri Hospital, Guri, Republic of Korea; SoVarGen Inc., Daejeon, Republic of Korea; Department of Otorhinolaryngology, Yonsei University College of Medicine, Seoul, Republic of Korea; Department of Neurosurgery, Yonsei University College of Medicine, Seoul, Republic of Korea; Center for Supercomputing Applications, National Institute of Supercomputing and Networking, Korea Institute of Science and Technology Information, Daejeon, Republic of Korea

## Abstract

Most somatic mutations arising during normal development present as low-level in single or multiple tissues depending on the developmental stage and affected organs^1-4^. However, it remains unclear how the human developmental stages or mutation-carrying organs affect somatic mutations’ features. Here, we performed a systemic and comprehensive analysis of low-level somatic mutations using deep whole-exome sequencing (WES; average read depth: ∼500×) of 498 multiple organ tissues with matched controls from 190 individuals. We found that early-stage mutations shared between multiple organs are lower in number but showed higher allele frequencies than late-stage mutations [0.54 vs. 5.83 variants per individual: 6.17% vs. 1.5% variant allele frequency (VAF)] along with less nonsynonymous mutations and lower functional impacts. Additionally, early- and late-stage mutations had unique mutational signatures distinct from tumor-originate mutations. Compared with early-stage mutations presenting a clock-like signature across all studied organs or tissues, late-stage mutations show organ, tissue, and cell-type specificity in mutation count, VAFs, and mutational signatures. In particular, analysis of brain somatic mutations shows bimodal occurrence and temporal-lobe-specific mutational signatures. These findings provide new insight into the features of somatic mosaicisms dependent on developmental stages and brain regions.

---

Somatic mutations persistently occur in normal cells during the entire human lifetime^1^. Although unaccompanied with unregulated proliferation, as seen in cancer, these somatic mutations often present a degree of clonality depending on time and origin. For example, variants in the early stages of development tend to affect multiple organs of different germ layers and show high variant allele frequencies (VAFs), whereas those in later stages localize with low VAFs^5,6^. Somatic variants that occur after birth are theoretically transient and restricted in a cellular level; however, mutations in stem or progenitor cells^7^ or variants that confer clonal expansion^8^ are persistent and accumulate during a lifetime and manifest a sufficient level of VAFs detectible in bulk-genome sequencing of tissues. Specifically, these tissue-level somatic mutations are crucial for the pathogenicity of non-cancerous or benign diseases, and the magnitude of aberrations is associated with their allele frequencies^9,10^. For example, mTOR-pathway-activating somatic mutations cause two types of intractable epilepsy (hemimegalencephaly and focal cortical dysplasia) depending on the time of mutation occurrence and VAFs (10–30% of VAFs in hemimegalencephaly, and 1–10% of VAFs in focal cortical dysplasia)^11-13^. Despite advances in the genetic identification of specific diseases, it still remains unclear how low-level but clone-forming somatic mosaicisms are generally characterized by the time and locations of their occurrence.

To address the questions, we performed a comprehensive analysis of low-level somatic mutations found in data from deep whole-exome sequencing (WES) of 498 tissues from 190 individuals (average read depth: ∼500×) **(Fig. 1a)**. The cohort consisted of multiple organs, including brain (*n*=301), blood (*n*=100), liver (*n*=60), heart (*n*=13), and other peripheral tissues (*n*=24). The 190 individuals included patients with ‘non-tumor’ neurological disorders (*n*=133), brain tumors (glioblastoma and ganglioglioma, *n*=19), and non-diseased controls (*n*=38) **(Supplementary Table 1)**. This cohort enabled multi-dimensional analysis and specifically a direct comparison with cancer mutations identified from a same analysis procedure.

**Figure 1.**
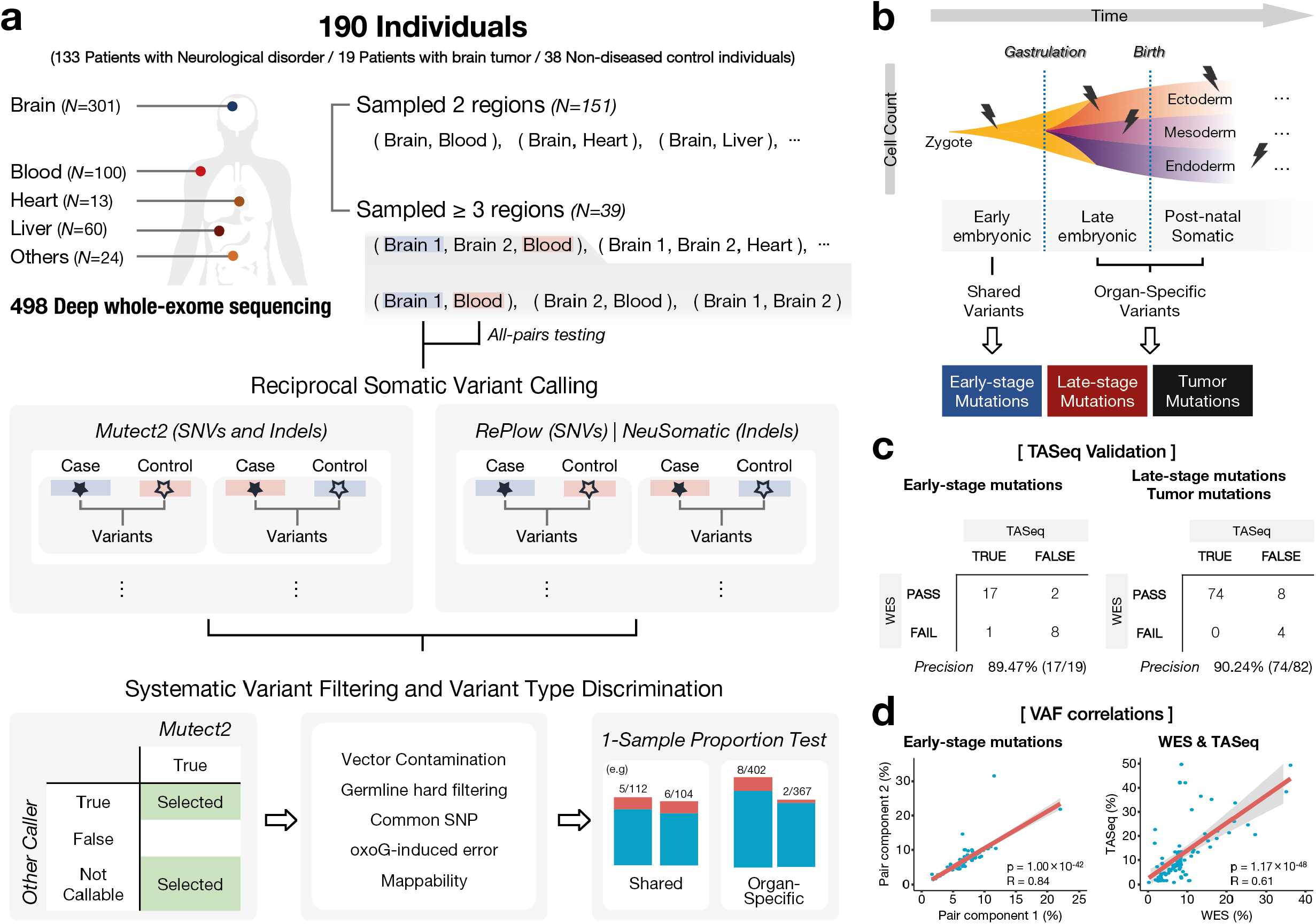
Detection of early- and late-stage somatic variants in brain and matched peripheral tissues. **a**, A schematic flow showing the bioinformatics pipelines of 301 brain tissues and 197 peripheral tissues from 190 individuals. To find somatic variants, Mutect2 and RePlow/NeuSomatic were used for reciprocal mutation calling by all-pairs testing, followed by post-call filtering. **b, c**, Early-stage, late-stage, and tumor mutations were classified with a highly accurate precision rate (89.47%, early-stage; and 90.24% in late-stage and tumor mutations). **d**, Correlation of VAFs from two matched tissues and WES and TASeq data. VAFs were highly concordant between paired tissues (*r* = 0.84; *P* < 0.0001) and WES and TASeq data (*r* = 0.61; *P* < 0.0001).

Regarding somatic mutations, we defined and used three different categories in the analysis: early-stage, late-stage, and tumor mutations **(Fig. 1b)**. Early-stage mutations were defined as mutations occurring during early embryonic development prior to gastrulation and shared in multiple-organs, whereas late-stage mutations included late embryonic (post-gastrulation) and post-natal somatic mutations restricted in a single organ. Based on the definition, somatic mutation calling was conducted using an ensemble of robust variant callers: Mutect2^14^, RePlow^15^, and NeuSomatic^16^ for 1,034 possible combinations of sample pairs. After strict filtration **(Fig. 1a)** and tests for organ specificity, we detected 103 early- and 997 late-stage mutations, as well as 583 tumor mutations. To validate the calls, 114 randomly selected single nucleotide variants (SNVs; ∼10% of non-tumor mutations) were sequenced by targeted amplicon sequencing (TASeq) to ultra-high depth (average: 507,856×) and Sanger sequencing. Our call set achieved high precision in both the early-stage (89.47%, 17/19) and late-stage and tumor mutations (90.24%, 74/82) **(Fig. 1c and Supplementary Table 2)**. High concordance in VAFs across tissues (Pearson’s correlation *r*=0.84; *P*=1.00×10^−42^) and between WES and TASeq data (*r*=0.61; *P*=1.17×10^−48^) confirmed the confidence of the calls **(Fig. 1d)**.

Additionally, we compared the quantitative traits of the mutations in terms of the number and allele frequency at different stages. On average, there were 0.54 early- and 5.83 late-stage somatic mutations per individual **(Fig. 2a)**. These numbers are roughly comparable to those of previous studies, which reported 0.53 shared and 3.15 non-shared somatic mutations in the brain (numbers were normalized to genomic size of 50 Mbp from whole-genome sequencing)^17,18^. It is possible that a slight increase in the number of late-stage (non-shared) might be due to the inclusion of blood samples, which are known to harbor ∼3-fold more mutations than other peripheral tissues^6^. Apparently, the numbers of mutations in normal tissues were substantially lower than those of tumors (30.00 per individual). The overall numbers of the late-stage and tumor mutations positively correlated with age (Pearson’s *r:* late-stage, 0.44; and tumor, 0.4) (**Fig. 2b)**. Conversely, we found no correlation between early-stage mutations and age, confirming that these mutations are well confined to the designated period. Regarding indels, 0.047 and 0.68 somatic indels were found in the early- and late-stage per individual, respectively **(Fig. 2a)**. The proportion of indels in the early-stage (∼8.7%) was slightly lower than that in the late-stage (∼11.7%) and tumors (∼10.7%). Because indels are more likely to be functionally damaging^19^, these results might represent lower tolerance to damaging mutations in the early developmental phase. On the other hand, VAFs of the mutations were higher in the early-stage (6.17 ± 3.32%) relative to the late-stage (1.50 ± 3.29%), which is consistent with the general expectation that somatic mutations that arise earlier present higher VAFs **(Fig. 2c)**. VAFs of early-stage somatic mutations have been measured in several studies with different criteria for inclusion and presented a diverse range (0.3–55%)^6,18,20^. Because none of the studies directly observed multi-organ-shared mutations using matched tissue sets from the same individuals, our analysis provides a more realistic distribution of VAFs for mutations occurring before gastrulation. Notably, VAFs of somatic indels in the early-stage were lower than those of somatic SNVs (indels vs. SNVs: 4.00% vs. 6.40%) but higher in the late-stage and tumors (2.75% vs. 1.34% in the late-stage; and 18.47% vs. 14.78% in tumors). The lower VAFs of indels, which represents lower cellular proportion and later occurrence, might be also associated with lower tolerance to damaging mutations in the early-developmental phase.

**Figure 2.**
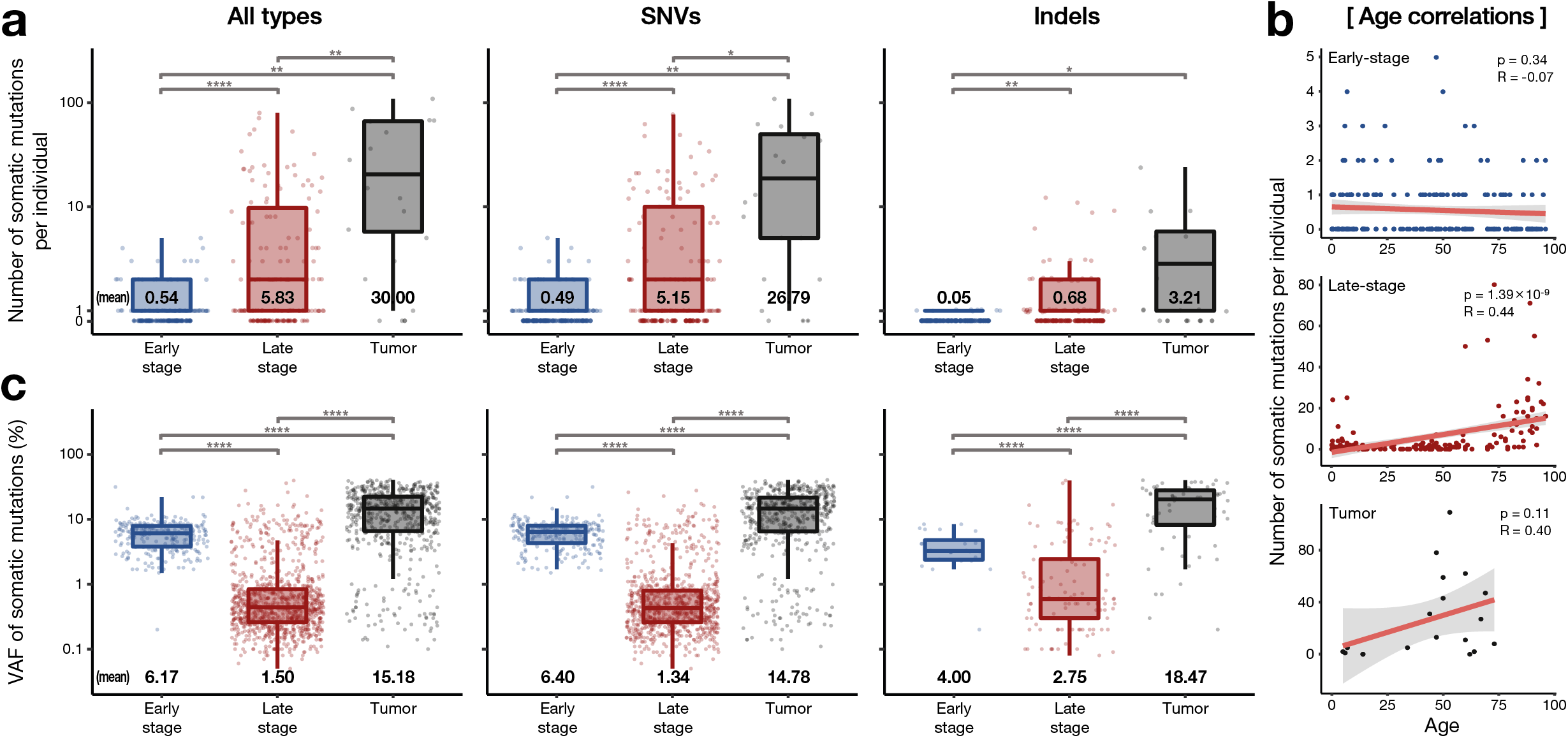
Basic descriptive statistics of somatic mutations. **a**, Number of somatic mutations per individual in early-stage, late-stage, and tumor mutation groups. **b**, Age correlation with somatic mutation counts in the groups. **c**, Average VAFs between the three mutation groups.

We then conducted mutation-profile analysis to investigate the underlying mutagenic processes **(Fig. 3a–d)**. *De novo* signature extraction of the 1,494 somatic SNVs (94 early-stage, 880 late-stage, and 520 tumor SNVs) identified three novel signatures **(Fig. 3a)**: signatures A, B1, and B2, all of which exhibited C>T as the major base substitution while showing additional T>C enrichment in signature A. Despite the overall similarity in mutational spectrum, especially between B1 and B2 (cosine similarity: 0.95), the clear distinction shown in the relative contribution to the sample groups confirmed the uniqueness of the signatures (*i*.*e*., signatures A, B1, and B2 dominantly contributed to the early-, late-stage, and tumor SNVs, respectively) **(Fig. 3b)**. This also implies that somatic mutations from different stages have distinguishing contexts. Mapping of the three signatures to COSMIC Mutational Signatures (v3.1; June 2020)^21^ identified clock-like SNV (SBS1, SBS5, and SBS40), and indel signatures (ID1, ID2, ID5, and ID8) as major components **(Fig. 3c)**. We noted that the relative contribution of the two well-known age-related signatures (SBS1 and SBS5) was altered from early- to late-stage SNVs (SBS1: 19% to 29%; and SBS5: 78% to 49%). The increased relative portion of SBS1 in late-stage somatic mutations appears to represent active proliferation and clonal expansion during late-embryonic and post-natal or aging periods^22,23^. Although the etiologies associated with most indel signatures remain unknown, the higher contributions of ID1 and ID2 in early-stage SNVs and ID5 and ID8 in late-stage SNVs were consistent with a previous finding^24^.

**Figure 3.**
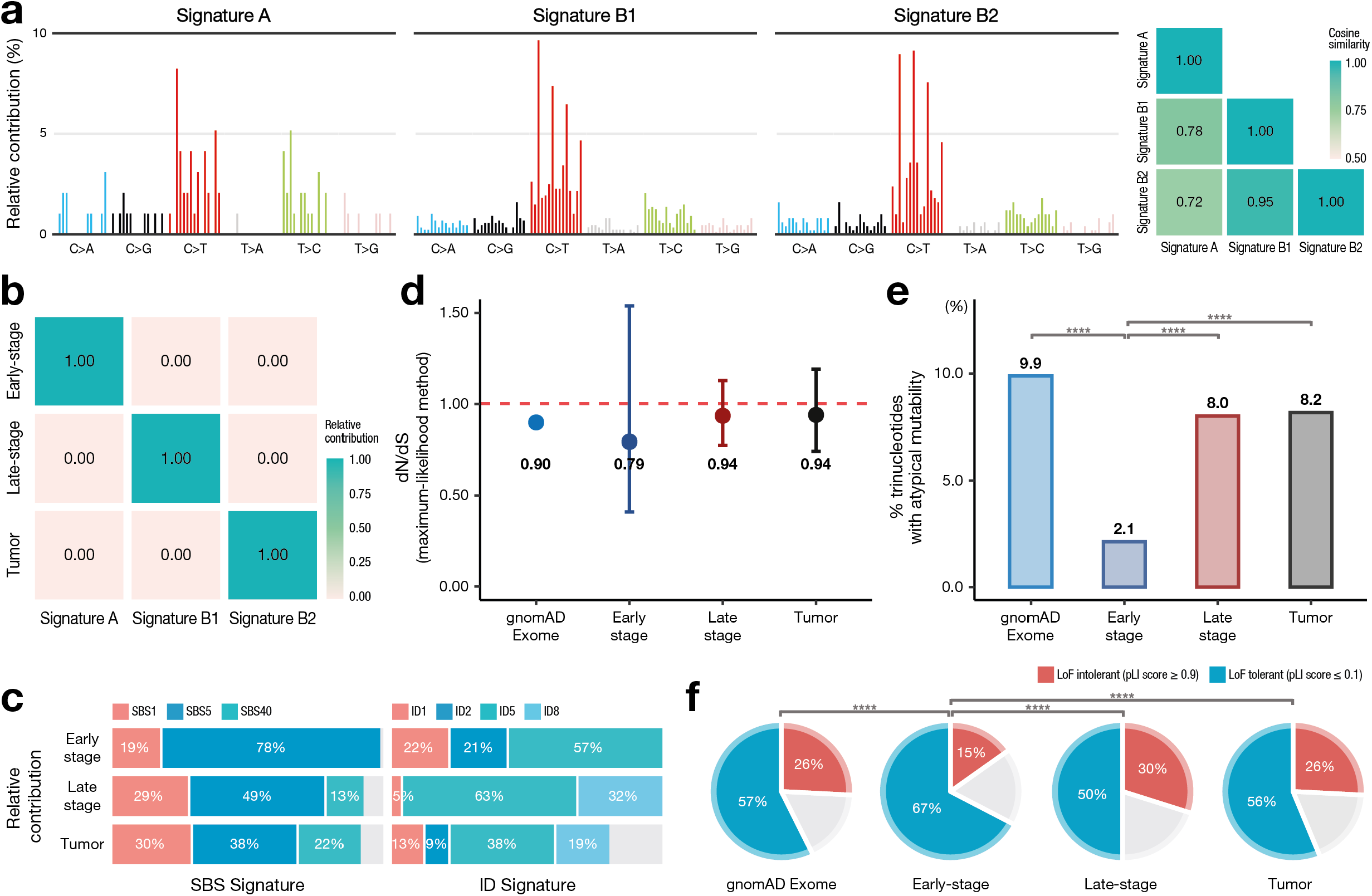
Mutational profile and functional analysis. **a**, *De novo* extraction of somatic mutations by non-negative matrix factorization. **b**, Each group was classified according to signature (A, B1, and B2). **c**, Relative contribution of the common clock-like signatures (SBS1, SBS5, and SBS40 for single-base substitutions and ID1, ID2, ID5, and ID8 for indels) from PCAWG signatures. **d**, The dN/dS score ratios, **e**, proportion of trinucleotides with atypical mutability, and **f**, pLI score for gnomAD Exome and each group.

Further assessment revealed differences between the early- and late-stage mutations in functional aspects. We found that early-stage mutations showed a lower ratio of non-synonymous to synonymous substitutions (dN/dS) (0.79) than did late-stage mutations (0.94), tumor mutations (0.94), and common germ-line coding variants (0.90; gnomAD Exome) **(Fig. 3d)**, indicating a stronger negative selection^25^. Additionally, early-stage mutations were less frequently (2.1%) located in trinucleotides with atypical mutability^26,27^ than were late-stage mutations (8.0%), tumor mutations (8.2%), and common germ-line coding variants (9.9%) **(Fig. 3e)**. Sites with atypical mutability are more highly mutated in cancer than is expected to occur randomly, indicating their functional significance and driverness in cancer^27,28^. Furthermore, genes that harbor early-stage mutations were lower in the probability of loss-of-function (LoF) intolerance (pLI score)^29^ **(Fig. 3f)**, indicating that early-stage mutations are more enriched in LoF-tolerant genes. These results collectively implied the strong selective pressure in the early embryonic stage^30,31^ that affects overall mutation characteristics that are less damaging possibly through the rejection of functionally-deleterious mutations.

We then investigated the characteristics of late-stage mutations, with a particular focus on diversity among organs and cell types. The numbers of mutations varied substantially by organ, with a smaller number in the brain (0.77 per individual) and higher number in the blood (9.24 per individual) relative to other peripheral organs (average: 1.13 per individual) **(Fig. 4a)**. However, the average number of VAFs was inversely proportional, with the highest number in the brain (7.32%) and the lowest in the blood (0.50%) **(Fig. 4b)**. Because VAFs generally decrease by the time of occurrence, we speculated that clonal somatic mutations in the brain occur relatively earlier but less frequently than those in the blood and other organs. The number of late-stage somatic mutations and the age of individuals showed a significant positive correlation (*r*=0.5; *p*=1.48×10^−6^) in only the blood **(Fig. 4c)**, which has been well-documented by post-natal clonal hematopoiesis^32,33^. Moreover, unsupervised hierarchical clustering of the three signatures (A, B1, and B2) of the late-stage mutations identified that those of the brain primarily comprise signatures A (early-stage) and B2 (tumor), whereas blood mutations are closer to signature B1 (late-stage) and B2 (tumor) **(Fig. 4d)**. These results suggest that late-stage somatic mutations in the brain present a bimodal-like occurrence during the embryonic period shortly after gastrulation and the post-natal period accompanied by a tumor-originating mutational signature.

**Figure 4.**
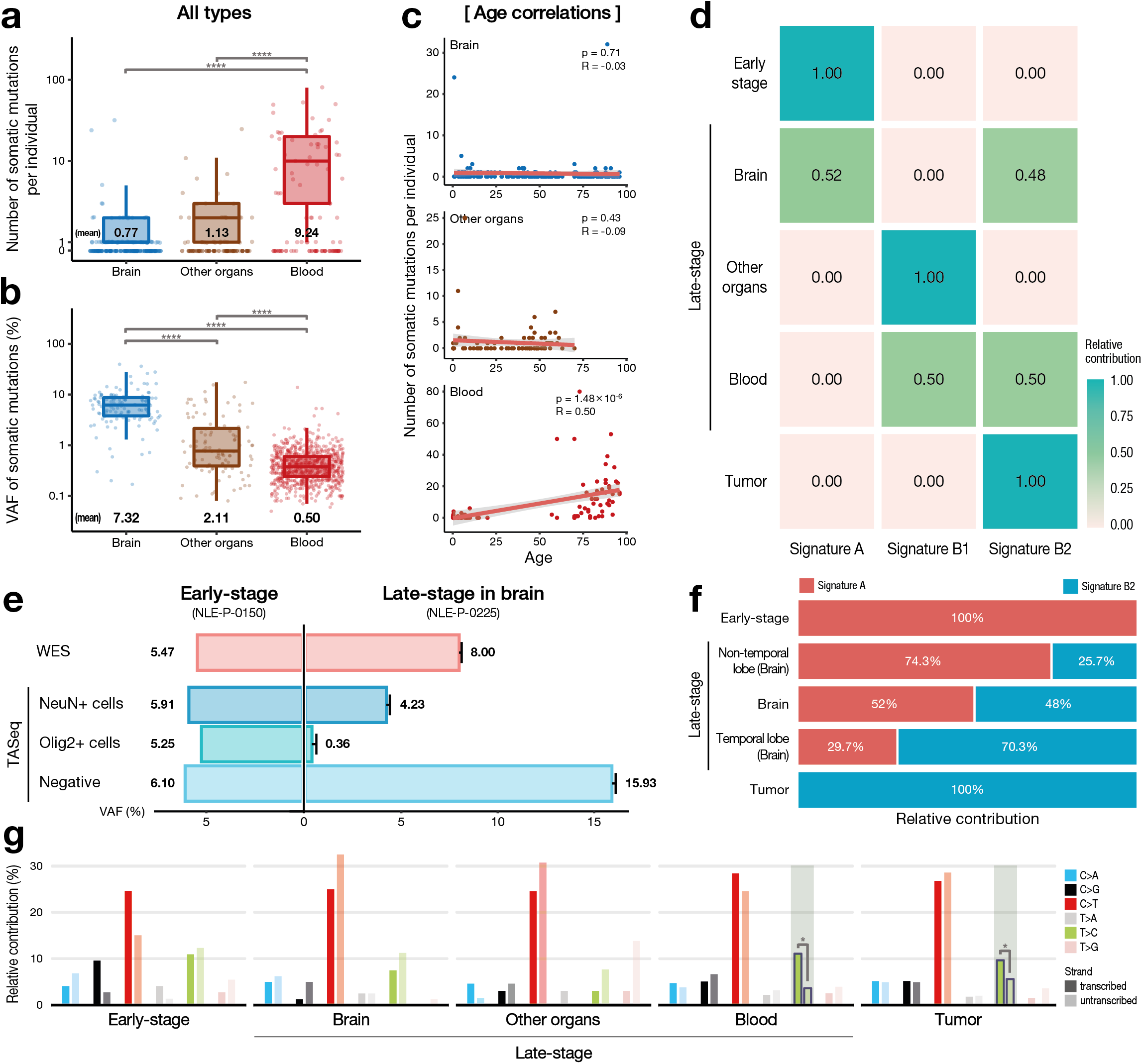
Analysis of late-stage mutations by mutation source for the organs (brain, blood, and other organs). **a, b**, Number of mutations per individual and VAF distribution. **c**, Age correlation with mutation counts. **d**, Unsupervised hierarchical clustering of late-stage mutations. Late brain somatic mutations were fit to signatures A and B2, whereas those in the blood were clustered to signatures B1 and B2. **e**, The VAFs of three different cell types [neuronal (NeuN+), oligogenic (Olig2+), and others (negative)] for early-stage and late-stage mutations in the brain. **f**, Signature distribution of late brain somatic variants divided among temporal and non-temporal areas or according to brain-disease status. **g**, Mutational-strand asymmetry. Late-onset blood and tumor mutations are noted as having strand-bias as T>C.

We then investigated the bimodal-like characteristics of the late-stage somatic mutations in the brain. First, we assessed the cell-type specificity of the somatic mutations in the brain by selecting two brain samples, which included one (NLE-P-0150) containing an early-stage mutation (5.47% VAFs) and the other (NLE-P-0225) five late-stage mutations (average: 8.00% VAF) **(Fig. 4e)**, each of which was sorted by fluorescence-activated nuclei sorting (FANS) to isolate three different cell types: neuronal (NeuN^+^), oligogenic (Olig2^+^), and others (negative). TASeq of the separated cell populations revealed that both early- and late-stage mutations are present in multiple cell lineages, but a large asymmetry of mutation frequencies among cell-types exists in the late-stage mutations **(Fig. 4e)**. These findings imply that the late-stage mutations in the brain occur later than the embryonic phase but relatively earlier during development in order to affect multiple lineages. We then subdivided the late-stage brain mutations into temporal and non-temporal areas and analyzed area-specific mutation signatures **(Fig. 4f)**. As previously reported, contributions to both areas were mainly from signatures A and B2; however, the degree of contribution of signature B2 was higher in the temporal lobe (70.3%) than non-temporal tissue (25.7%), revealing that the characteristics of somatic mutations in the temporal lobe are closer to those of tumor mutations. We speculated that the tumor-like mutational signatures in the temporal lobe might originate from neurogenesis activity (e.g., dentate gyrus) that confers clonal proliferation, as reported previously^34^. Furthermore, the strand specificity of the late-stage mutations in blood and tumor mutations showed enrichment of T>C mutations on transcribed strands **(Fig. 4g)**. Because transcription coupled repair occurs more frequently with higher transcription levels and this bias is increased in actively replicating templates^35,36^, we again confirmed that clonal expansion-derived somatic mutations were included in the blood, similar to those in tumors^37^.

In summary, based on a large scale of deep whole exome sequencing data using a total of 498 matched sample pairs from multiple organs in 190 individuals, we provided a more detailed picture of low-level but clone-forming somatic mutations, the counts, and characteristics of which are distinguished by time and space. We found that early-stage mutations, which arise prior to gastrulation and are shared in multiple organs, are lower in number and have lower functional impact than late-stage mutations restricted within a single organ. Moreover, we showed that late-stage mutations are associated with human mutational processes in the late-embryonic and post-natal developmental stages but that vary by organ, tissue, and cell lineages. In particular, late-stage mutations in the brain showed a bimodal-like occurrence over developmental stages and asymmetry of mutational features across brain-cell types and regions. Regarding the asymmetry of somatic mutations, asymmetric cell divisions resulting from early cellular bottlenecks of stochastic clonal selection contributed to an uneven variant fraction according to developmental timing^6,38^. These findings suggest that the VAFs of clone-forming somatic mutations reflect not only the timing of the mutation but also the cell fitness and cell-type specificity for given somatic mutations. Overall, the well-defined characteristics of each mutation group and target tissue according to their developmental period can confer an accurate representation of currently-observable somatic mutations and a better understanding of how they were generated.

## Methods

### Patient samples

The acquired freshly frozen brain and peripheral samples of 24 autism spectrum disorder (ASD) and five non-ASD cases from the National Institute of Child Health & Human Development (Bethesda, MD, USA) included various brain regions, such as the frontal, temporal, occipital, and cerebellar areas. Paired samples with other organs were derived from 13 ASD cases and five non-ASD case, and brain samples were obtained from 11 ASD cases. The Stanley Medical Research Institute (Rockville, MD, USA) supplied genomic DNA of brain tissue and other matched organs for 25 non-schizophrenia and 26 schizophrenia cases. Additionally, the Stanley Medical Research Institute provided genomic DNA for brain and matched liver tissues from patients with major depressive disorders. Fresh frozen brain samples of Alzheimer’s disease (AD) were provided from the Netherlands Brain Bank (project number Lee-835) for 96 brain and matched blood samples for AD and non-demented control cases, and 15 samples of AD and non-demented control cases were obtained from the Human Brain and Spinal Fluid Resource Center (West Los Angeles Healthcare Center, Los Angeles, CA, USA), which is sponsored by NINDS/NIMH (Bethesda, MD, USA), the National Multiple Sclerosis Society (Raleigh, NC, USA), and the US Department of Veterans Affairs (Bethesda, MD, USA). Fresh frozen samples of lumbosacral lipoma were supplied from the Severance Children’s Hospital of Yonsei University College of Medicine (Seoul, Republic of Korea). Bone tissues of non-syndromic craniosynostosis patients were provided from the Severance Hospital of Yonsei University College of Medicine. Subjects with refractory epilepsy, including focal cortical dysplasia and non-lesional epilepsy, and who had undergone epilepsy surgery were enrolled through the Severance Children’s Hospital of Yonsei University College of Medicine. Subjects with glioblastoma and ganglioglioma were enrolled from the Severance Hospital of Yonsei University College of Medicine and satisfied diagnostic criteria according to the 2016 World Health Organization Classification of Tumors of the Central Nervous System^39^. We were provided freshly-frozen samples of resected brain lesions.

### Deep WES

Genomic DNA was extracted with either the QIAamp mini DNA kit (Qiagen, Hilden, Germany) from freshly frozen brain tissues or the Wizard genomic DNA purification kit (Promega, Madison, WI, USA) from blood according to manufacturer instructions. Each sample was prepared according to Agilent library preparation protocols (Human All Exon 50 Mb kit; Agilent Technologies, Santa Clara, CA, USA). Libraries were subjected to paired-end sequencing on an Illumina Hiseq 2000 and 2500 instrument (Illumina, San Diego, CA, USA) according to the manufacturer’s instructions) with confidence-mapping quality (mapping quality score ≥ 20; base quality score ≥ 20).

### Data processing and systematic variant calling

We checked the quality of the raw sequencing reads using FastQC^40^ (v.0.11.7) software. The FASTQ-formatted sequencing reads of each sample that passed the quality check were aligned to the human reference genome (build 38; NCBI, Bethesda, MD, USA) using the BWA-MEM^41^ algorithm and converted into a BAM file. The initial BAM file was updated with read groups, and duplicate information was excluded as it progressed through the steps using Picard^42^ and GATK^43^. Additionally, we performed local realignment and base-quality recalibration with GATK tools for each exome. BAM files that successfully underwent all of these steps were then used to measure contamination between samples, with the probability of swapping assessed using NGSCheckMate^44^ software and cross-contamination tested using GATK tools. Vecuum^45^ software was used to check for vector contamination during library construction, and Depth of Coverage (GATK) was used to measure sequencing depths. All processes not described in detail were performed based on the GATK best-practice pipeline.

Two or more tissue samples from each individual were paired using all-pairs testing. We performed the somatic mutation-detection pipeline (paired mode) with sample pairs as inputs using a three somatic variant caller; Mutect2^14^ somatic variant-calling pipeline, excluding the panel of normal creation (SNVs and Indels), RePlow^15^ (SNVs), and NeuSomatic^16^ with the control of the false detection rate control performed by Varlociraptor^46^ (Indels).

All mutations meeting the following conditions were removed from the initial mutation-detection results in the VCF format: oxoG-induced errors according to the method described by Costello et al.^47^, common single-nucleotide polymorphisms by NCBI dbSNP^48^ (build 153), segmental duplication and simple repeat regions according to the UCSC database^49^, a mappability score >0.8 by Umap^50^, and presence of an off-target region^51^ whole genome without exome and the untranslated region.

### Decisions regarding early and late mutations

After the removal of artefacts, somatic mutations with “PASS” results for both Mutect2 and other caller filters (RePlow/NeuSomatic) were classified as late-stage mutations. If the source of the sample was related to a brain tumor, it was separately regarded as a tumor mutation.

Early-stage mutations were initially categorized as such if the filter result of Mutect2 was “normal artifact” or RePlow (for only SNVs) was “normalFilter,” respectively. Additionally, these were assigned this category if they were called in Mutect2 only but not in RePlow. After confirming amino acid changes and genomic location, to confirm that the same mutation was detected from each individual, the validity of the mutation was statistically verified using the one-sample proportion test. The VAFs of each mutation were used as a criterion to determine whether the ratio of the ‘ref’ and ‘alt’ alleles of the other mutations satisfied the null hypothesis. Common mutations in different samples from each individual were tested, and mutations satisfying the criteria were classified as early-stage mutations.

### Validation sequencing of candidate mutations using deep-targeted amplicon sequencing or Sanger sequencing

We then performed validation sequencing by randomly selecting mutations from each group. For validation, we used deep-targeted amplicon sequencing or Sanger sequencing of PCR-amplified DNA. Primers for PCR amplification were designed using Primer3 software (http://bioinfo.ut.ee/primer3-0.4.0/)^52^. Target regions were amplified by PCR using specific primer sets and high-fidelity PrimeSTAR GXL DNA polymerase (Takara, Shiga, Japan). Sanger sequencing was performed using BigDye Terminator reactions and loaded onto a 3730xl DNA analyzer (Applied Biosystems, San Francisco, CA, USA).

### Bioinformatics analysis

All somatic mutations excluded false positives by validation sequencing were annotated using VEP^53^ (v.99.0) with “-everything -plugin ExACpLI” options. The results were evaluated using an in-house script to analyze the descriptive statistics of the properties of the basic mutations, effect of each gene, and possible correlations with patient demographics (age, disease, etc.). Non-negative matrix factorization-based novel signature extraction (200 iterations) and transcriptional strand-bias analysis were performed using the MutationalPatterns program^54^. The signature and 96-types variant contexts were fitted to clockwise Pan-Cancer Analysis of Whole Genomes (PCAWG) single-base substitution and small insertions and deletions signatures by deconstructSigs^55^, Mutalisk^56^ (date of use: March 2020), and YAPSA^57^. Mutability was calculated using NCBI MutaGene^26,27^ (v.0.9.1.0) distributed as a Python package. The maximum-likelihood dN/dS method was applied by dNdScv (Wellcome Sanger Institute, Cambridge, UK)^25^.

### Nuclei extraction and FANS

Frozen brain samples were minced using pre-chilled razor blades and one or two drops of lysis buffer [0.2% Triton X-100, 1× protease inhibitor, and 1 mM DTT in 2% bovine serum albumin (BSA) in phosphate-buffered saline]. Lysis buffer (1 mL) was added to the homogenate and mixed by pipetting, after which the lysate was fixed in 1% paraformaldehyde at room temperature for 10 min, and the fixed lysate was quenched with 0.125 M glycine at room temperature for 5 min. The homogenate was then washed with suspension buffer (1 mM EDTA and 2% BSA) and filtered with 40-µM cell strainer. The sample was then incubated with anti-NeuN (mature neuronal marker; 1:1000) and anti-Olig2 (oligodendrocyte lineage marker; 1:500) overnight at 4°C, followed by washing with suspension buffer and staining with the secondary antibody for 1 h at 4°C. After washing with suspension buffer, nuclei were passed through a 40-µM cell strainer and stained with 1 µg 4′,6-diamidino-2-phenylindole. Nuclei used to isolate each cell type were analyzed and sorted using a MoFlo Astrios EQ cell sorter (Beckman Coulter, Brea, CA, USA). Nuclei pellets were centrifuged for 10 min at 1500*g* and processed immediately for gDNA extraction using a QIAamp DNA micro kit (Qiagen) according to manufacturer instructions.

## Acknowledgements

This research was supported by the National Research Foundation of Korea (NRF) grant funded by the Korea government (MSIT) (No. 2019R1A2C2008050 to S. K.), the Suh Kyungbae Foundation (to J.H.L.), and a National Research Foundation of Korea (NRF) grant funded by the Korean Ministry of Science and Information and Communication Technology (ICT) (No. 2019R1A3B2066619 to J.H.L)

We thank the Netherlands Brain Bank (Lee-835) for Alzheimer and unaffected control cases., the National Institute of Child Health & Human Development for providing Autism and unaffected control cases, the Stanley Medical Research Institute for the brain and peripheral DNA provided of schizophrenia, major depressive disorders, and unaffected control cases, Seoul National University Hospital, Seoul National University College of Medicine for providing lumbosacral lipoma and non-syndromic craniosynostosis, and Severance Hospital, Yonsei University College of Medicine for providing samples of brain tumor and epilepsy, which were supplied to J.H.L.

## Author contributions

S.K. and J.H.L designed and initiate the study. H.S. and J.H.K conducted main analysis. H.S. devised analysis pipeline and performed bioinformatics analysis. J.H.K. worked on sample organization, validation sequencing, and FANS. I.B.K, M-H.K., and N.S.S. prepped human tissue samples and performed whole-exome sequencing. D-S.K. performed the epilepsy surgeries, collected patient samples, and managed patient information. J.L. worked on analysis of sequencing data. H.S., J.H.K., J.H.L., and S.K. worked on data interpretation, and wrote the manuscript with input from coauthors. J.H.L and S.K. led the project.

## Competing interests

J.H.L. is co-founder and CTO of SoVarGen Inc., which seeks to develop new diagnostics and therapeutics for brain disorders. The other authors declare no competing interests.

